# A coarse-grained model for disordered proteins under crowded conditions

**DOI:** 10.1101/2025.03.01.640997

**Authors:** Arriën Symon Rauh, Giulio Tesei, Kresten Lindorff-Larsen

**Author notes:** **For correspondence:** (KLL).

## Abstract

Macromolecular crowding may strongly affect the dynamics and function of proteins, with intrinsically disordered proteins being particularly sensitive to their crowded environment. To understand the influences of crowding on chain compaction and phase separation behaviour of disordered proteins, both experiments with synthetic crowders—like polyethylene glycol (PEG) and ficoll—and theoretical models and molecular simulation approaches have been applied. Here, we developed a residue-based coarse-grained model for PEG that is compatible with the protein CALVADOS model. To achieve this, we optimised model parameters by comparing simulations with experimental data on single-chain PEG and on PEG-induced compaction of disordered proteins. With our model we show how titrations of PEG can be used to quantify phase separation propensities of proteins that are not prone to phase separate strongly. We illustrate this for both variants of the low-complexity domain of hnRNPA1, and for wild-type and a redesigned variant of *α*-Synuclein. Notably, we observe that the PEG crowding response changes between charge patterning variants of *α*-Synuclein, which is not the case for the variants that vary the number the of aromatic residues in the low-complexity domain of hnRNPA1. We expect that our model will be useful for the interpretation of crowding experiments with disordered proteins, and we envisage it to be a starting point for in-silico explorations of proteins with weak propensities to phase separate.

## Introduction

In contrast to the conditions of most biochemical experiments, the natural environment of a protein in a cell is densely packed with macromolecules. Since up to 25% of the cellular volume can be occupied by biomolecules, the effective volume available for any single molecule in the cell is markedly reduced compared to typical in-vitro experimental conditions (***Zimmerman and Trach, 1991***; ***Ellis and Minton, 2006***; ***Zhou et al., 2008***; ***Boersma et al., 2015***; ***Rivas and Minton, 2016***; ***Alfano et al., 2024***). This macromolecular crowding gives rise to entropic excluded volume effects, such as depletion interactions, as well as ‘weak’ enthalpic interactions due to the close proximity of molecules. Together, these interactions can lead to altered protein diffusion, stability, interactions, complex formation, and overall conformational dynamics (***Berg, 1990***; ***Hong and Gierasch, 2010***; ***Mikaelsson et al., 2013***; ***Guin and Gruebele, 2019***; ***Zosel et al., 2020***; ***Cubuk and Soranno, 2022***). Thus, crowding can, for example, modify aggregation and binding kinetics, and influence complex protein behaviour in processes like phase separation (***Atha and Ingham, 1981***; ***Haire et al., 1984***; ***Annunziata et al., 2002***; ***Li et al., 2013***; ***André and Spruijt, 2020***; ***Chauhan et al., 2024***). Studying and modelling proteins in crowded environments provides insights into how the local environment modulates protein function through excluded volume effects and transient interactions.

While all proteins are susceptible to crowding, intrinsically disordered proteins and regions (IDPs hereafter) are perhaps particularly prone to perturbations from dense environments, as they are fully or partially unfolded under physiological conditions and adopt highly heterogeneous conformational ensembles (***Cubuk and Soranno, 2022***). IDPs play a wide range of roles in biology, and their conformational properties can be linked to functions, for example in signalling and gene regulation (***Wright and Dyson, 2015***; ***Moses et al., 2023***; ***Holehouse and Kragelund, 2023***).

Experimental methods such as nuclear magnetic resonance (NMR), electron paramagnetic resonance (EPR) and single-molecule fluorescence spectroscopy (smFRET) have been used to study the effects of crowding, for example, by studying proteins in solutions of synthetic crowders like polyethylene glycol (PEG), dextran and ficoll (***Sarkar et al., 2013***; ***Köhn and Kovermann, 2020***; ***Pierro et al., 2024***; ***Alfano et al., 2024***). Such experiments have shown that both the crowder (mass) concentration and molecular weight (MW) affect the compaction of IDPs (***Soranno et al., 2014***; ***Stringer et al., 2023***). Excluded volume effects can to some extent be described by scaled-particle theory, which models crowders as hard-spheres (***Reiss et al., 1959***). Further theoretical developments have improved the description of the effect of crowders on unfolded proteins by representing the protein volume with a Gaussian distribution, which allows for overlap between crowder particles and polypeptide chain (***Minton, 2005***). However, for flexible polymers such as PEG, the crowding effect on IDPs is influenced by the degree of polymerisation, which modulates the balance between the relative contribution of volume exclusion and crowder–protein interactions (***Soranno et al., 2014***). Therefore, crowding can be modelled more accurately by taking both the polymeric nature of the protein and the crowder into account using a Flory-Huggins-based theory for polymer solutions (***Soranno et al., 2014***). This framework bridges the enthalpic effects for lower-MW crowders with the increasingly dominant entropic effects for higher-MW crowders.

Crowding also affects the demixing of protein solutions, which is one of the potential mechanisms for the formation of biomolecular condensates (***Banani et al., 2017***; ***Mittag and Pappu, 2022***; ***Pappu et al., 2023***). Since some IDPs may exert their biological functions by driving or modulating the formation of biomolecular condensates, understanding how crowding affects the phase separation (PS) of IDPs is crucial. The conditions under which a protein phase separates are sensitive to temperature, pH, ion concentrations, and the concentrations of other macromolecular components (***Banani et al., 2017***; ***Alberti, 2017***; ***Dignon et al., 2020***; ***Pappu et al., 2023***). Thus, while the propensity of a protein to phase separate is encoded in its sequence (***Wang et al., 2018***; ***Martin et al., 2020***; ***von Bülow et al., 2024***), this propensity may be modulated by both entropic excluded volume effects and the weak interactions between crowders and phase separating proteins (***Atha and Ingham, 1981***; ***Qian et al., 2022, 2024***; ***Chauhan et al., 2024***). Examples include using PEG to induce the PS of proteins such as *α*-Synuclein, Tau, NPM1, YAP, MED1, and BRD4 (***Sabari et al., 2018***; ***Cai et al., 2019***; ***Ray et al., 2020***; ***Kanaan et al., 2020***; ***André et al., 2023***; ***Swain et al., 2024***).

Molecular simulations have been used to complement experiments and theory to describe the molecular effects of crowding including on phase separation (***Miller et al., 2016***; ***Heo et al., 2022***; ***Mathur et al., 2024***; ***Grassmann et al., 2024***; ***Chauhan et al., 2024***). For example, dissipative particle dynamics simulations have been used to study crowding effects on a FUS-based condensate system (***Shillcock et al., 2023***). Lattice-based simulations and determination of virial coefficients have also been applied to study crowding by simulations, and to support extrapolations of the propensity to phase separate (quantified by the saturation concentration, *C*_sat_) from crowder titrations (***Edmond and Ogston, 1968, 1970***; ***Chauhan et al., 2024***). We and others have used off-lattice, one-bead-per-residue models for IDPs to study phase separation driven by IDPs (***Dignon et al., 2018a***; ***Dannenhoffer-Lafage and Best, 2021***; ***Regy et al., 2021a***; ***Tesei et al., 2021***; ***Joseph et al., 2021***; ***Latham and Zhang, 2021***; ***Jung et al., 2024***; ***Jussupow et al., 2024***). These models enable efficient computational studies of IDPs and their PS behaviour by representing each amino acid with a single bead described with a bead size, *σ*, and a hydropathy (‘stickiness’) parameter, *λ*. In the case of our previously described CALVADOS model, the *λ* parameters were determined by optimisation against single-chain compaction data from SAXS and PRE experiments (***Tesei et al., 2021***; ***Tesei and Lindorff-Larsen, 2022***). PEG-driven crowding of a protein represented by the CALVADOS coarse-grained model has been studied using a hard-sphere representation of PEG (***Mizutani et al., 2023***). To gain a more detailed description of crowding or PEG-ylation it may be advantageous to represent the crowder using a more detailed model, such as the PEG model in the popular coarsegrained molecular dynamics model Martini (***Lee et al., 2009***; ***Rossi et al., 2012***; ***Grünewald et al., 2018***).

To enable the study and analysis of the effects of crowding on IDPs and their PS, we have developed a residue-based coarse-grained model for PEG that is compatible with the protein CALVADOS model. We parameterise the model using measurements of compaction of PEG and the effect of PEG on the compaction of IDPs. Similar to previous work studying how phase separation depends on crowder concentrations (***Chauhan et al., 2024***) and how denaturant titrations have been used to study protein stability (***Lindorff-Larsen and Teilum, 2021***) we show how crowder titrations can be used to quantify even weak propensities to phase separate for variants of both the low-complexity domain of hnRNPA1 (A1-LCD) and for *α*-Synuclein (*α*-Syn).

## Results

### Capturing PEG chain dimensions across molecular weights

To enable modelling the effects of macromolecular crowding on IDPs, we aimed to develop a coarse-grained model of PEG that is compatible with our CALVADOS model for simulations of IDPs (***Tesei et al., 2021***; ***Tesei and Lindorff-Larsen, 2022***). In the CALVADOS model, each amino acid is represented by a single ‘bead’ whose characteristics are described by three parameters, a size parameter (*σ*), a ‘stickiness’ or hydrophobicity parameter (*λ*), and—for titratable residues—a charge. CALVADOS has been parameterised using a top-down strategy (***Norgaard et al., 2008***) in which we optimised the *λ* values against experimental NMR and small-angle X-ray scattering data on IDPs (***Tesei et al., 2021***).

We began by examining possible values of *λ* and *σ* for describing solutions of PEG and then (see further below) examined interactions between PEG and proteins. Specifically, we first scanned parameters for hydrophobicity (*λ*_PEG_) and size of the monomer unit (*σ*_PEG_) for reproducing experimentally measured chain dimensions in single-chain simulations of PEG with MWs ranging from 400 Da to 30 kDa (Fig. 1B, for an overview of all plots see Fig. S1). In our choice for the range of values to scan for *σ*_PEG_, we mainly restricted ourselves to values close to or below the size of a glycine bead in CALVADOS (*σ*_Gly_=0.45 nm, ***Kim and Hummer*** (***2008***)) as it is the closest in size and atomic composition to a PEG monomer (C_2_H_5_NO_2_ vs. C_2_H_6_O).

**Figure 1.**
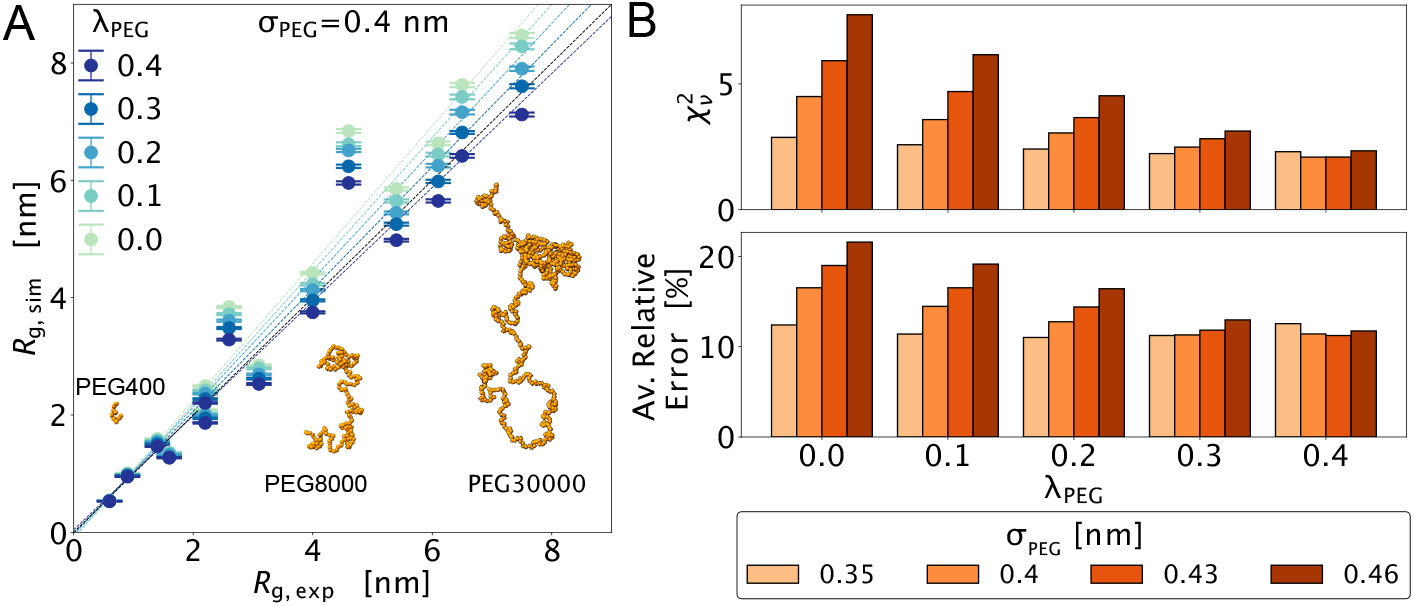
Scanning parameters for reproduction of single-chain properties of linear PEG chains with different molecular weights. (A) Agreement between simulations and experimental scattering data of PEG chain dimensions for different combinations of *λ*_PEG_ and *σ*_PEG_. For an illustration of the range of molecular weights we scan, we add snapshots of PEG400, PEG8000 and PEG30000 with 9, 181 and 681 monomers, respectively. (B) Comparison between experimental and calculated ***R***_g_ in simulations of PEG with *σ*_PEG_ = 0.4 nm and varying *λ*_PEG_ values. In the top panel, we show the values of the reduced 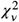 and in the bottom panel the relative error.

In the scan, we observe two trends for the agreement with the ***R***_g,exp_ values determined from scattering experiments (***Sherck et al., 2020***), which we quantify using 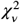 and the average relative error (Fig. 1A). First, when we increase the values of *λ*_PEG_ from 0.0 to 0.3 the agreement increases; however, it plateaus around *λ*_PEG_=0.4. Second, with lower values of *σ*_PEG_, the effect of varying *λ*_PEG_ becomes less discernible (Fig.1A). Based on these results we select *σ*_PEG_=0.4 nm as a balance between a physically plausible size of the PEG monomer and the ability to capture PEG chain dimensions. We note that this value is also close to that of the ‘small’ bead used in the Martini model to represent PEO/PEG (*σ*_PEO, martini 3_=0.41 nm) (***Grünewald et al., 2018***).

### Probing PEG induced compaction of intrinsically disordered proteins

Our goal was to develop a model that captures the balance between PEG–PEG and protein–PEG interactions. We therefore fixed *σ*_PEG_ = 0.4 nm and scanned *λ*_PEG_ in the range from 0.0 to 0.3 while probing the interactions between PEG and IDPs. To minimise the complexity of the model, we use a single value of *λ*_PEG_ for both PEG–PEG and protein–PEG interactions, while describing protein– protein interactions with the amino-acid-specific *λ* parameters from the CALVADOS 2 model (***Tesei and Lindorff-Larsen, 2022***).

We thus compared simulations with previously determined values of the end-to-end distance (***R***_ee_) from single molecule Förster resonance energy transfer (smFRET) measurements (***Soranno et al., 2014***). Specifically, we compared simulations to experimental smFRET data for activator for thyroid hormones and retinoid receptors (ACTR) and the N-terminal domain of HIV-1 integrase (IN) in the presence of PEG400 and PEG8000 at volume fractions (*ϕ*_PEG_) ranging from 0% to just below 40% (***Soranno et al., 2014***), as illustrated in figure 2D). Initially, we also considered experimental data on the highly charged C-terminal part of prothymosin-*α*, but decided against using them to tune *λ*_PEG_ since this might inadvertently absorb deficiencies in the Debye-Hückel electrostatic model into the PEG model. The results for ACTR with both PEG400 (Fig. 2A) and PEG8000 (Fig. 2B) show that the best agreement with experiments is achieved with *λ*_PEG_ around 0.2–0.3. We aggregated the results from both PEG400 and PEG8000 across several values of *ϕ*_PEG_ in a single *χ*^2^ value and repeated these calculations also for IN (Fig. S2). In both cases, we find that *λ*_PEG_ = 0.2 gives rise to the best overall agreement with experiments (Fig. 2C).

**Figure 2.**
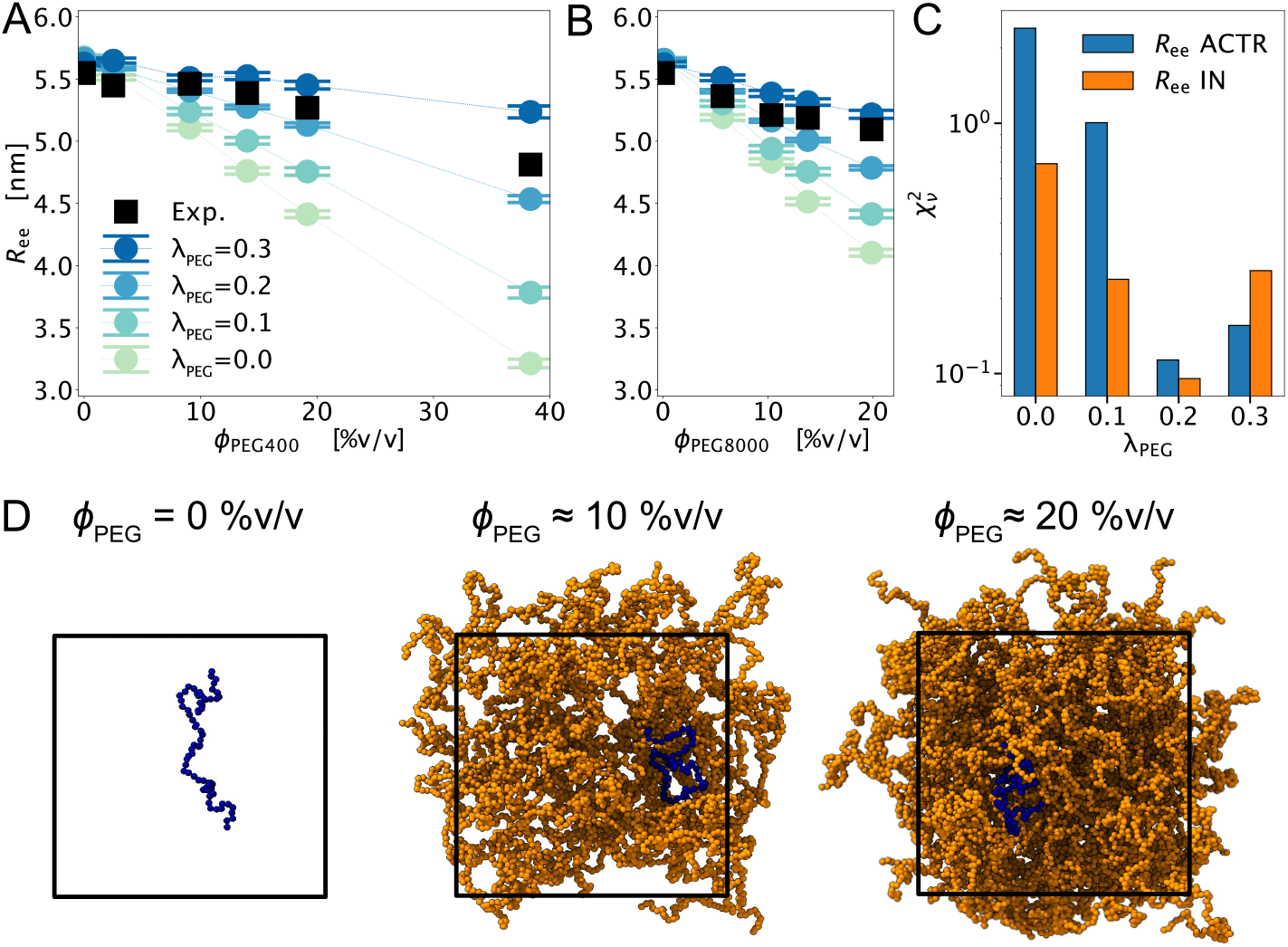
Tuning *λ*_PEG_ by comparison with PEG-induced compaction of IDPs. End-to-end distances (***R***_ee_) of ACTR with varying volume fractions (*ϕ*_PEG_) of (A) PEG400 (degree of polymerization, DP = 9) (B) and PEG8000 (DP = 181) simulated with different values of *λ*_PEG_. Error bars indicate the standard error of the mean estimated through blocking analysis. Experimental ***R***_g_ values (***Soranno et al., 2014***) were converted to ***R***_ee_ values and shown as black squares. (C) Reduced *χ*^2^ between experimental and simulated values of ***R***_ee_ of thyroid hormones and retinoid receptors (ACTR, blue) and the N-terminal domain of HIV-1 integrase (IN, orange) indicate best agreement with *λ*_PEG_ = 0.2. (D) Representative snapshots from simulations of a single chain of ACTR (dark blue) with volume fractions of 0% (1:0), 10% (1:36) and 20% (1:69) PEG8000 (orange chains) illustrate how crowded the protein environment is at these volume fractions.

### Phase separation of hnRNPA1 LCD is linearly dependent on PEG concentrations

Having determined that the combination of *σ*_PEG_ = 0.4 nm and *λ*_PEG_ = 0.2 provides a reasonable balance between PEG–PEG and protein–PEG, we proceeded to study how crowding affects the propensity of IDRs to phase separate. In this context, phase separation refers to the spontaneous demixing of a solution into a protein-rich ‘dense phase’ and a dilute phase (***Pappu et al., 2023***). Phase separation occurs when the concentration of the protein is above the so-called saturation concentration (*C*_sat_); *C*_sat_ depends both on the sequence of the protein and the conditions, and is roughly determined by the balance between protein–protein and protein–solvent interactions. The resulting driving force for phase separation is often quantified by *C*_sat_, or by the transfer free energy between the protein-dilute and -dense phase, *ΔG*_transfer_ = *RT* ln *C*_dil_ *C*_den_, where *C*_dil_ = *C*_sat_ is the dilute phase concentration when phase separation occurs.

We have recently estimated that ca. 5% of naturally occurring human disordered regions will undergo homotypic phase separation with *ΔG*_transfer_ *<* −5 kJ/mol at ambient conditions (***von Bülow et al., 2024***), meaning that most disordered sequences will not easily undergo spontaneous phase separation. This in turn makes it difficult to estimate—both experimentally and computationally— the propensity of these proteins to phase separate. Experimentally, one commonly applied approach to induce phase separation of proteins with a weaker driving force is to add crowding agents such as PEG (***Annunziata et al., 2002***; ***Sabari et al., 2018***; ***Cai et al., 2019***; ***Ray et al., 2020***; ***Kanaan et al., 2020***; ***André et al., 2023***). In some cases, this approach has been used to study phase separation of different proteins, (***Sabari et al., 2018***; ***Kanaan et al., 2020***; ***Hochmair et al., 2022***; ***Jo et al., 2022***), with interpretations sometimes under the implicit assumption that their relative propensities to phase separate are mostly unaffected by the presence of PEG.

An alternative approach is to use crowder titrations and extrapolate the measurements to the absence of crowder (***Edmond and Ogston, 1968, 1970***; ***Chauhan et al., 2024***); this approach is analogous to the commonly applied approach to study the thermodynamic stability of proteins by unfolding them by addition of denaturant and extrapolating the results to the absence of denaturant (***Lindorff-Larsen and Teilum, 2021***). A log-linear response has previously been found and used to study protein solubility/saturation concentration as a function of PEG concentration (%w/v) (***Edmond and Ogston, 1968***; ***Atha and Ingham, 1981***; ***Annunziata et al., 2002***; ***Shillcock et al., 2023***; ***Chauhan et al., 2024***).

To examine this log-linear relationship in more detail we performed simulations of the phase separation of the low-complexity domain of hnRNPA1 (A1-LCD) in the presence of increasing concentrations (0 to 10 %w/v) of both PEG400 and PEG8000, similarly to previous work (***Chauhan et al., 2024***). Our choice of A1-LCD as a model system is motivated by the fact that its phase separation behaviour is well studied (***Martin et al., 2020, 2021***; ***Bremer et al., 2022***) and that the CALVADOS 2 model describes the propensity of A1-LCD and various sequence variants well (***Tesei and Lindorff-Larsen, 2022***).

In simulations of A1-LCD in a ‘slab’ geometry, we find that both PEG400 and PEG8000 increase the driving force for phase separation (making *ΔG*_transfer_ more negative) and we observe a linear response in *ΔG*_transfer_ with an increasing PEG concentration (Fig. 3A; for dilute and dense phase concentrations see Fig. S3). To quantify both the specific PEG response (slope) and estimate the *ΔG*_transfer_ in the absence of PEG (the -intercept), we fit *ΔG*_transfer_ from simulations with PEG concentrations between ca. 2%w/v–10%w/v. The results are well described by this linear fit apart from at the highest concentration of PEG, which is most likely due to difficulties in converging simulations in these very crowded systems. As expected, the linear extrapolations appear to converge to the same value in the absence of PEG, and indeed an F-test shows that a fit to the two curves with a common intercept is not distinguishable from one with two separate intercepts (*p* = 0.33). We also performed simulations in the absence of PEG and find that the value extrapolated from the crowder titration is within error of the value obtained directly from simulations of A1-LCD without PEG (Fig. 3A). We also find that the two different crowders give rise to distinct responses in the propensity of A1-LCD to phase separate, with PEG8000 having a substantially larger (ca. 60%) effect than PEG400 at the same volume fractions (Fig. 3A), in line with previous observations of distinct effects of differently sized crowders (***Atha and Ingham, 1981***; ***Soranno et al., 2014***; ***Chauhan et al., 2024***).

**Figure 3.**
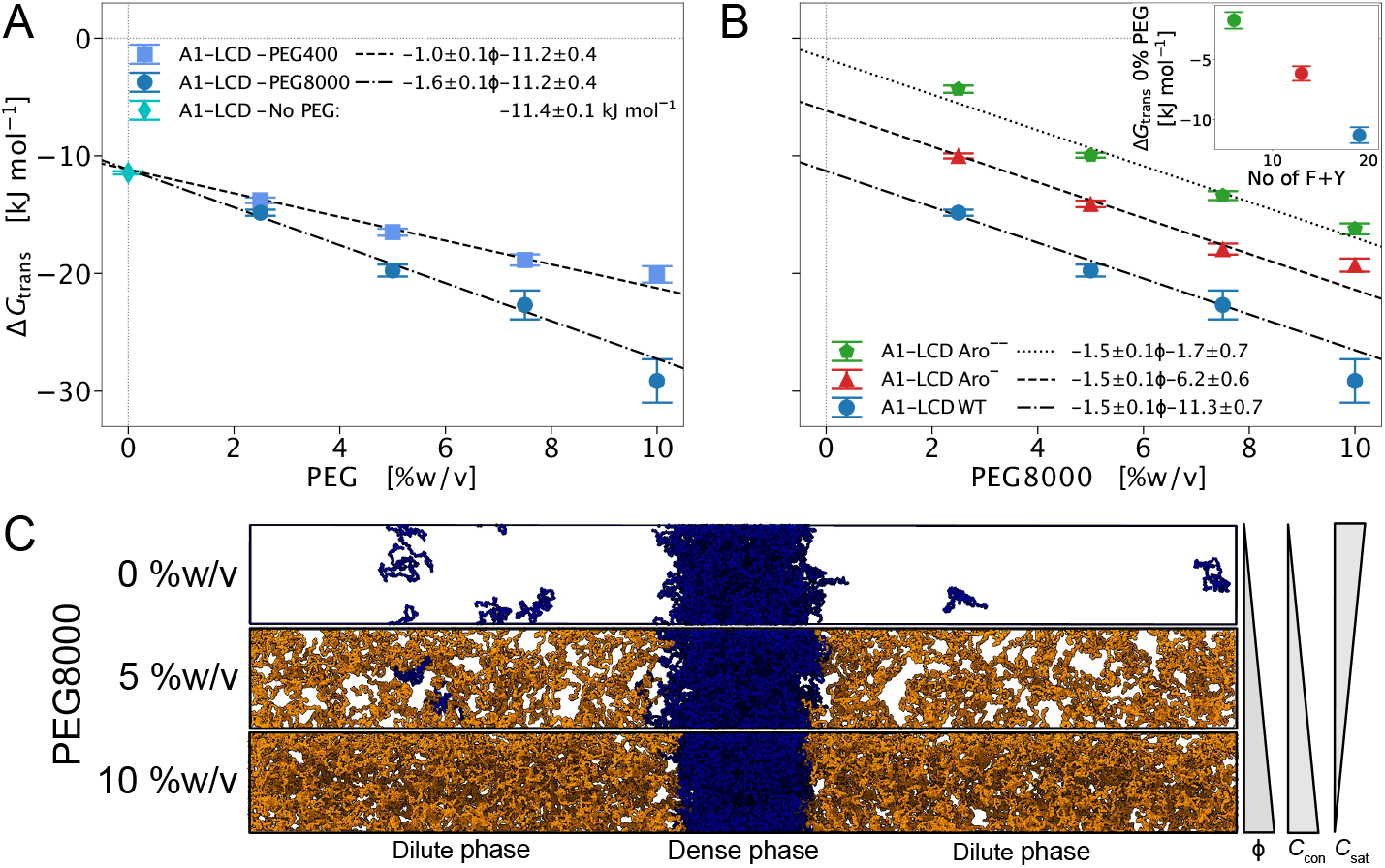
The phase separation propensity of WT and aromatic variants of the hnRNPA1-LCD responds linearly to PEG concentration. (A) A1-LCD PS propensity (*ΔG*_transfer_) with both PEG400 (squares) and PEG8000 (circles). The linear fits were performed with the *y*-intercept as a shared global parameter (3-parameter fit), which is not a significantly worse fit in comparison with the 4-parameter fit (F-test *p* = 0.33). The A1-LCD PS propensity calculated from replicate slab simulations without PEG is shown at 0%w/v (diamond). (B) PS propensities of WT A1-LCD and the aromatic variants Aro^−^ (red triangles) and Aro^−−^ (green pentagrams) show a similar response (slopes) to crowding by PEG8000. Here the linear fits were performed with a shared slope (4-parameter fit), which does not perform significantly worse than a 6-parameter fit (F-test *p* = 0.22). (C) Representative snapshots from slab simulations with 100 chains A1 (dark blue) with w/v-percentages of 0%, 5% (100:127) and 10% (100:254) PEG8000 (orange chains, 181 monomers) illustrating how PEG both lowers the *C*_sat_ with increasing concentrations and also increases the effective protein concentration in the condensate (*C*_con_).

Further exploring the differences related to crowder size, we find that up to 10% of the total amount of PEG400 chains at any stage of the titration mixes in with the protein in the condensed phase, whereas PEG8000 chains more clearly segregate out of the protein-dense phase with increasing PEG concentration. Thus, in this case, we find a trend of the larger PEG polymers partitioning less, which could be an entropic effect as the cost of admitting a chain into the condensate likely increases with its degree of polymerization. A similar trend has been seen in PS experiments of bovine *γ*D-crystallin (***Annunziata et al., 2002***).

### Linear extrapolation captures the differences in intrinsic PS propensity of hnRNPA1-LCD aromatic variants

We have now established that the PEG model produces an apparent linear response in PS propensity (*ΔG*_transfer_) with increasing PEG concentration, and showed that this allows us to distinguish between the effects of crowders of different sizes and correctly extrapolate the intrinsic phase propensity of A1-LCD in the absence of PEG. To further explore these extrapolation capabilities, we turned to two well-described aromatic variants of A1-LCD where a third (Aro^−^) and two-thirds (Aro^−−^) of the aromatic residues have been mutated (***Martin et al., 2020***). These variants illustrate the relationship between A1-LCD PS propensities and the aromatic content, where a decrease in aromatic content comes with a decrease in PS propensity. In the case of A1-LCD Aro^−−^ variant, the decrease in aromatic content results in PS behaviour that is not easily measurable in experiments, thus for example necessitating the use of extrapolation from a PEG titration to quantify the PS propensity of this variant. To see if we can reproduce the relationship between the aromatic content and PS behaviour with extrapolations, we performed simulations of Aro^−^ and Aro^−−^ with increasing concentrations of PEG8000 (0 to 10 %w/v).

From our ‘slab’ geometry simulations of the aromatic variants, we observe a linear response in *ΔG*_transfer_ as a function of the PEG concentration with similar slopes but different -intercepts (Fig. 3B, the separate saturation and condensate concentrations are plotted in Fig. S4). To quantify the specific PEG response (slope) and estimate the *ΔG*_transfer_ in the absence of PEG (the -intercept) for A1-LCD WT and aromatic variants, we fitted linear curves with a globally optimised shared slope, assuming that the variants and WT protein show a similar crowding response to PEG8000. This global fit, with a shared slope for the three curves (4-parameter fit), does not perform significantly worse than a fit with separate slopes (*p* = 0.22). We note that the use of a shared slope here is analogous to using a common *m*-value in analysing protein stability data (***Kellis Jr et al., 1989***; ***Lindorff-Larsen, 2019***). The shared slope for the A1-LCD variants with PEG8000 is also consistent with the previously fitted slope for A1-LCD WT with PEG8000 (−1.5 ± 0.1 vs −1.6 ± 0.1, Fig. 3A & B). Extrapolation of the fitted linear curves indicates that, without PEG, A1-LCD aromatic variants Aro^−^ and Aro^−−^ have *ΔG*_transfer_ of −6.2 ± 0.6 and −1.7 ± 0.7 kJ mol^−1^, respectively. When we examine these values as a function of aromatic content (number of phenylalanine and tyrosine residues; Fig. 3B), we see a linear trend of decreased PS propensity with decreasing aromatic content. This apparent linear trend in *ΔG*_transfer_ without PEG is in agreement with the results of critical temperatures from binodals determined in a study combining experiments and lattice-based simulations (***Martin et al., 2020***). Moreover, we demonstrate here that our model is applicable for the determination of the intrinsic PS propensity of a protein that shows no phase separation without the addition of PEG, through linear extrapolation of a titration curve. For illustrative purposes we also show snapshots of the the phase separated protein and the crowder (Fig. 3C).

### Crowding-induced phase separation of Ddx4

Having demonstrated the capabilities of the CALVADOS PEG model for extrapolating PS trends of A1-LCD WT and aromatic variants, we wanted to explore a separate system. We set out to compare our simulation results with estimates of the saturation concentration (*C*_sat_) from a set of ‘Capillary flow experiments’ (Capflex) of an N-terminal construct of the DEAD-box Helicase 4 protein (Ddx4n1) in the presence of increasing PEG3000 concentrations (2-7 %w/v) (Table S3, ***Stender et al***. (***2021***)). To achieve this, we performed slab simulations of Ddx4n1 with PEG3000 in a range between 2 and 7 %w/v. From the initial simulations, it was clear that, for PEG3000 concentrations above 3 %w/v, the sampled *C*_sat_ values are noisy due to difficulties in sampling. If a protein is prone to phase separate more strongly, *C*_sat_ is low, which means that sampling a chain in the protein-dilute phase will be harder and artefacts arise more easily. When adding PEG to the simulation this problem is exacerbated as the crowder increases the stability of the condensate. So, to enable sampling of a less noisy titration curve, we decided to add three data points around 1 %w/v (Fig. 4A). The simulations display a log-linear PEG response in the range between 0.7 and 2 %w/v, which we can subsequently quantify by fitting an exponential function. We did this for both the experimental and the simulated *C*_sat_ titration series to facilitate the comparison between PEG responses. It is clear that the overall crowding trend (constant in the exponent), as determined with the Capflex method, is overestimated in the simulations with our PEG model. The extrapolated *C*_sat_ without PEG from our simulations (95 ± 6 µM) only slightly overestimates the extrapolated value from the PEG3000 crowding experiments (71±2 µM), however, it is within error of the average *C*_sat_ measured in the absence of PEG (100 ± 2 µM, Table S4) (***Stender et al., 2021***). What we show here for Ddx4n1 is that even though our model is not fully quantitatively accurate on the crowding response in a PEG3000 titration, it accurately predicts *C*_sat_ in the absence of PEG by linear extrapolation of the titration curve.

**Figure 4.**
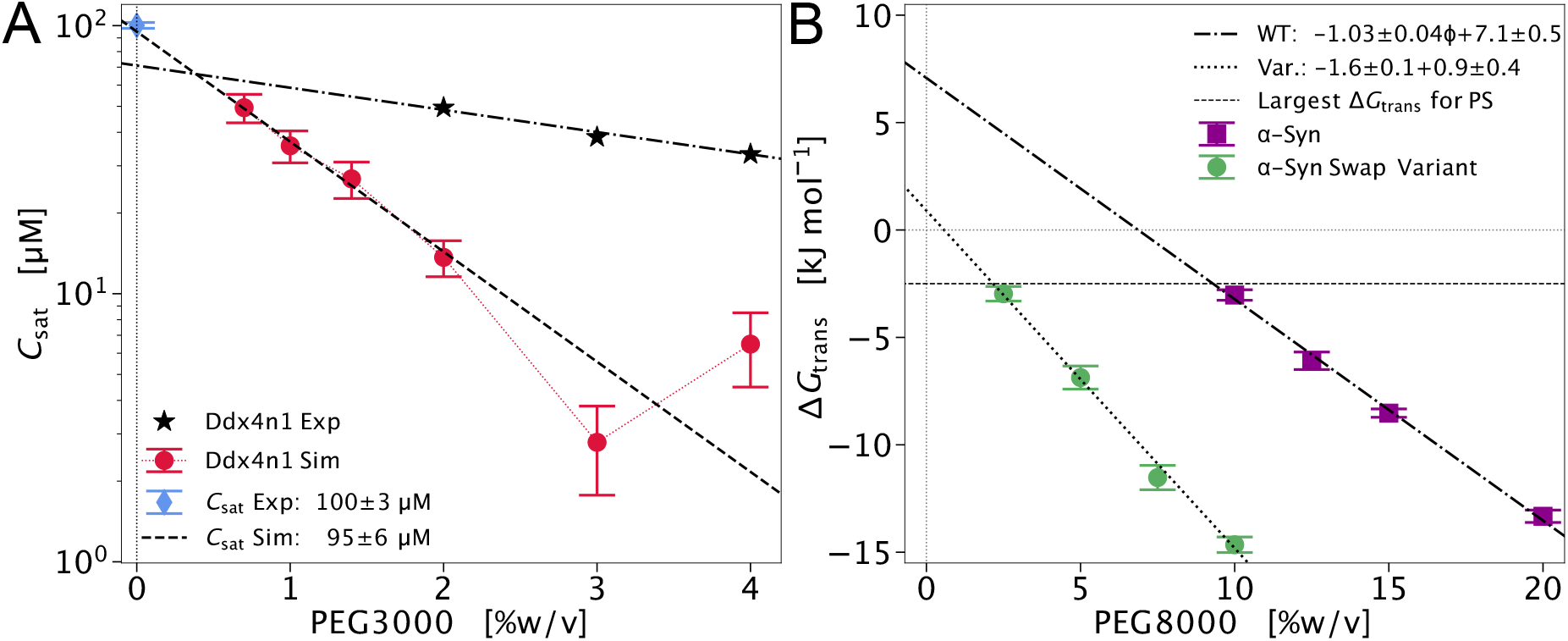
Validation and application of the PEG model. (A) Comparison between simulated and experimental PEG3000 crowder titrations for Ddx4n1. *C*_sat_ values are from co-existence simulations (red dots) and Capflex measurements with PEG (black stars) and without PEG (blue diamond). All experimental data is taken from ***Stender et al***. (***2021***). (B) Simulated PEG8000 titration curves for wild-type (WT) *α*-Syn (dark magenta squares) and a charge-segregated swap variant (green dots) optimised for higher chain compaction. The linear fit was performed with no shared parameters (4-parameter fit), which performs significantly better than a 3-parameter fit (F-test *p* = 0.01).

### Phase Separation of *α*-Synuclein and a Charge-segregated Swap Variant

Finally, we showcase how our PEG model can be applied to study the phase separation behaviour of a weakly PS-prone protein that does not show PS in a standard CALVADOS simulation. As a test system, we chose the Parkinson’s disease-related protein *α*-Syn, which requires the addition of crowders for homotypic PS to occur in in-vitro experiments at physiologically relevant conditions (***Ray et al., 2020, 2023***). As a first step, we inspect whether our model aligns with experimental trends on the PS of *α*-Syn induced by PEG. For this, we performed slab simulations of *α*-Syn with 0 to 20 %w/v PEG8000, covering the ranges explored in fluorescence experiments (***Ray et al., 2020***; ***Sawner et al., 2021***). Our slab simulations indicate that *α*-Syn phase separates with 10 to 20 %w/v PEG8000, with an overall linear response to crowder concentration (Fig. 4B, dark magenta squares) whereas, below 10 %w/v PEG8000, *α*-Syn does not exhibit PS (Fig. S7). At 10 %w/v PEG8000, the *ΔG*_transfer_ value is around −2.5 kJ mol^−1^. These results are in qualitative agreement with the experimental trends described by ***Sawner et al***. (***2021***) and ***Ray et al***. (***2020***), where *α*-Syn also shows no PS for PEG8000 concentrations below 10%w/v, at pH 7.4 and for protein concentrations between 100 and 200 µM. To explore the PS behaviour of *α*-Syn further, we quantified the crowding response and determined an estimate of the *ΔG*_transfer_ without PEG by extrapolation of a linear fit in the range between 10 and 20 %w/v PEG8000. The extrapolation suggests that the *ΔG*_transfer_ without PEG is 7.1 ± 0.5 kJ mol^−1^ for *α*-Syn (Fig. 4B in dash-dotted line). A positive *ΔG*_transfer_ is in line with the absence of PS without the addition of PEG in experiments. This number can for example be compared compared to *ΔG*_transfer_ for the similarly sized WT A1-LCD (−11.3 ± 0.7 kJ mol^−1^) or the more weakly phase separating Aro^−−^ variant (−1.7 ± 0.7 kJ mol^−1^) (Fig. 3B).

Next, we decided to use the PEG titration approach to examine the PS propensity of *α*-Syn in more detail. For homopolymers and a number of IDPs it has been shown that the propensity for homotypic PS is correlated with chain compaction, as the intra-molecular interactions that allow a chain to be compact also give an indication of how favourable the inter-molecular interactions are in a mixture of multiple chains of the same protein (***Lin and Chan, 2017***; ***Dignon et al., 2018a***; ***Martin et al., 2020***; ***Bremer et al., 2022***; ***von Bülow et al., 2024***). Based on this symmetry, we can hypothesise that a modification of the sequence that makes the single chain more compact could also imply an increased PS propensity. To explore this, we analysed a variant of *α*-Syn designed to be more compact than the wild-type protein while maintaining the same amino acid composition (***Pesce et al., 2024***). Simulations of this designed variant show that it has an R_g_ approximately 1 nm lower than that of WT *α*-Syn (Fig. S5). In sequence space, this was mainly achieved by increasing the segregation of charged residues through consecutive positional swaps between pairs of residues. We quantify charge patterning using the sequence descriptor *κ* (***Das and Pappu, 2013***), which takes values close to zero for highly charge-segregated sequences and values close to 1 for sequences with well-mixed charges. The change in, *κ* between the compact swap variant and WT *α*-Syn is considerable and amounts to *Δκ*_wt→var_ = −0.423 (see Table S2 for an overview of sequence descriptors and Fig. S7 for a visual inspection of the sequences).

To examine whether the symmetry between single-chain and PS behaviour also holds true here, so that the more compact swap variant would show an increased PS propensity, we also performed a titration series of slab simulations with increasing PEG8000 concentrations for the swap variant. The addition of PEG in the simulations of the swap variant is needed just as with WT *α*-Syn, as it also does not show phase separation in a standard CALVADOS simulation. Our simulations show that the swap variant exhibits PS above 2.5 %w/v PEG8000 and its *C*_sat_ at 10%w/v is lower than that of WT *α*-Syn at 20%w/v (Fig. 4B and Fig. S6 in green dots). This indicates that the PS propensity for the compact variant is stronger relative to WT *α*-Syn. To quantify this increased PS propensity, we performed a linear fit for the *α*-Syn swap variant between 2.5 and 10%w/v PEG8000 and determined the *ΔG*_transfer_ without PEG through extrapolation. *ΔG*_transfer_ without PEG is 0.9 ± 0.4 kJ mol^−1^, which gives a 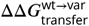 of 6.2 ± 0.6 kJ mol^−1^. Our simulations, thus, indicate that charge segregation with minimal perturbation of hydropathy patterning not only increases compaction but also the PS propensity for *α*-Syn. We note that the PS propensity of the *α*-Syn variant (0.9 ± 0.4 kJ mol^−1^) is of comparable magnitude as for the weakly phase separating Aro^−−^ variant (−1.7 ± 0.7 kJ mol^−1^).

From our simulation data, we also observe a different response to crowding between WT *α*-Syn and its swap variant with perturbed charge patterning. Specifically, we find that the slopes of the linear fits of the titration curves, 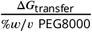, differ by a factor of ≈ 1.6, with the slope for the variant being considerably steeper. This difference in slope is significant since fitting the curves with linear functions with distinct slopes outperforms a fit with a shared slope (*p* = 0.01).

This observation is in contrast with our findings for PEG titrations of the aromatic variants of A1-LCD, where the PEG-crowding response is not significantly different between the variants. Together, these observations show that the PEG response (slope) may sometimes, but not always, depend on the sequence of the protein, similarly to the effect of crowding of the compaction of individual proteins (***Miller et al., 2016***). This in turns means that one should be cautious when inferring and quantifying differences in intrinsic PS behaviour between variants (in the absence of crowder) by only comparing data at a single PEG concentration, as this implicitly assumes that the crowder affects each variant in the same way.

## Discussion

To aid the interpretations of crowding experiments for IDPs, we here implemented a residue-level model for the crowding agent PEG, compatible with the CG MD CALVADOS protein model. We did this through a data-driven parameterisation of a bead representing a PEG monomer. In our parameterisation strategy, we explore both PEG–PEG interactions, by examining the compaction of PEG chains with varying MWs, and PEG–protein interactions, by comparing with crowder-induced compaction experiments. Subsequently, we show how, in a similar manner to previous studies (***Chauhan et al., 2024***), this model can be applied in an in-silico titration experiment for explorations of homotypic phase separation propensities of IDRs, leveraging the previously described log-linear relationship between PS and macromolecular crowding (***Atha and Ingham, 1981***; ***Haire et al., 1984***; ***Annunziata et al., 2002***; ***Li et al., 2013***). With the titration approach, we demonstrate that in an exploration of aromatic variant effects for A1-LCD, we can reproduce the trends of decreased phase separation propensities with decreasing aromatic content (***Martin et al., 2020***). Further, for the first time with the CALVADOS model, we explore the PS behaviour of *α*-Syn, a protein with a weak propensity to phase separate, and observe a PS trend that is qualitatively in line with experimental data (***Ray et al., 2020***; ***Sawner et al., 2021***).

A shortcoming of this work is that we cannot demonstrate the quantitative accuracy of the model in reproducing the PEG response of PS, as our predictions of the PEG titration curves for Ddx4n1 deviate from the corresponding experimental data (***Stender et al., 2021***). However, we show that we can extrapolate to the experimental *C*_sat_ value for Ddx4n1 in the absence of PEG, which was obtained from Capflex measurements under similar conditions as for the PEG titration data. In future work, it would be useful to compare experiments and simulations of phase separation across a broader range of proteins and PEG sizes.

On a practical note, when simulating systems with a weak PS propensity, a point of attention is to critically assess what constitutes a quantifiable dense phase and thus determine the lowest *ϕ*_PEG_ value at which PS is induced. In our simulations of both WT and variants of A1-LCD and *α*-Syn, we observe the formation of a protein-rich slab for values of *ΔG*_transfer_ below -2.5 kJ mol^−1^. On the other hand, when a system exhibits a highly stable condensate, sampling an accurate value for *C*_sat_ is challenging since exchange events between the coexisting phases become rare. This problem worsens in systems at high PEG concentrations, where *C*_sat_ is low and the sampling efficiency deteriorates due to the increased number of particles in the simulation and the longer decorrelation times. For example, on an NVIDIA Tesla V100 GPU we can sample ≈ 4.5 µs/day for 100 copies of A1-LCD without PEG whereas the simulation speed decreases to ≈ 1.5 µs/day and ≈ 0.7 µs/day with 10% and 20%w/v PEG8000, respectively. Moreover, to achieve convergence on *C*_sat_ values, we need to double the simulation time from ≈ 10 µs, which is generally sufficient in the absence of PEG, to at least ≈ 20 µs at high PEG concentrations.

We applied the model to gain insights into the PS of IDPs induced by PEG of different MWs. Both low-MW (400 Da) and high-MW (8000 Da) PEG partition outside the protein-dense phase. However, shorter PEG chains are less excluded at high PEG concentrations, which is likely due to entropic effects being weaker for shorter chains and therefore less dominant over enthalpic protein–PEG interactions (***Annunziata et al., 2002***; ***Stewart et al., 2023***). We also examined how PEG influences the PS of IDPs of different sequence patterning. Notably, titration curves for *α*-Syn variants with different charge patterning display different slopes, whereas aromatic variants of A1-LCD do not. This suggests that when comparing PS behaviour between two protein variants, a full titration should be performed, as measurements at a single PEG concentration are insufficient to estimate the change in PS propensity in the absence of PEG. Similarly to the use of *m*-values in analysing protein stability data (***Shortle, 1995***; ***Geierhaas et al., 2007***), we envisage that there is useful information also encoded in the analysis of the PEG response, and future studies could examine the relationship between protein sequence, PEG size and the slope.

In summary, we have implemented a PEG crowder model by optimising using experimental data taking both PEG–PEG and PEG–protein interactions into account. Our results support and extend previous work that shows how PEG titrations can aid in determining the intrinsic propensities of a protein to phase separate (***Annunziata et al., 2002***; ***Chauhan et al., 2024***), and illustrate how to apply them for the study of proteins with low PS propensities. With an efficient molecular model of PEG, we can now study the specific polymer behaviour of PEG, for example, under a transition from a dilute to a semi-dilute regime (***Cohen et al., 2009***; ***Palit et al., 2017***). Furthermore, we anticipate that this model, in combination with the crowder titration approach, will be a valuable tool for future in-silico investigations of weakly phase-separating proteins and their variants, facilitating a more detailed understanding of the sequence determinants governing ‘weaker’ PS. With this we can, for instance, investigate how the PS of *α*-Syn relates to its aggregation behaviour, by analysing variants known to show increased aggregation (***Koo et al., 2008***; ***Larsen et al., 2025***). More generally, we can also expand the training set of a recently developed PS predictor (***von Bülow et al., 2024***) to enhance its coverage of protein sequences with lower PS propensities, which should provide improved efficiency and boost large-scale explorations of PS-related variant effects. Another promising direction is the integration of our model with the recently published CALVADOS 3 framework for multi-domain proteins (***Cao et al., 2024***), which allows us to investigate crowding effects across a broader range of systems, including full-length proteins, for which more additional experimental data is available (***Stringer et al., 2023***; ***Qian et al., 2022***; ***Jo et al., 2022***; ***Kaur et al., 2019***).

## Methods

### Simulations

We performed all coarse-grained molecular dynamics simulations with openMM (v7.5) (***Eastman et al., 2017***) in the NVT ensemble, using a Langevin integrator with a time step of 10 fs and a friction coefficient of 0.01 ps^−1^. We model IDRs using CALVADOS 2, a molecular model where each residue is represented as a single bead located at the position of the C *α* atom. The hydrophobicity ‘stickiness’ scale of this HPS-type model (***Dignon et al., 2018b***) was determined through a Bayesian parameter-learning procedure to capture the sequence-dependent compaction of IDRs (***Tesei et al., 2021***; ***Tesei and Lindorff-Larsen, 2022***). We model bonds with a harmonic potential,

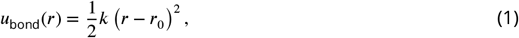

where *k* = 8030 kJ mol^−1^ nm^−2^, *r*_0_ = 0.38 nm for amino acids and *r*_0_ = 0.33 for PEG beads. We model the non-bonded ionic interactions with salt-screened electrostatic interactions using a Debye-Hückel potential truncated and shifted at a cutoff of 4 nm,

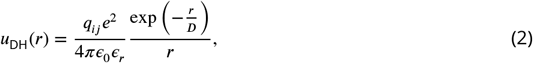

where *q*_*ij*_ is the product of the charge numbers for interacting residue beads, *e* is the elementary charge, *e*_0_ is the vacuum permittivity, and the Debye-length *D* of an electrolyte solution with an ionic strength 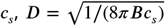, where *B*(*ϵ*_*r*_) is the Bjerrum length. We model the temperature-dependent dielectric constant of the implicit aqueous solution *ϵ*_*r*_ with an empirical relationship (***Akerlof and Oshry, 1950***),

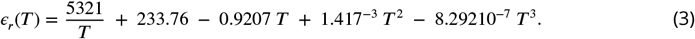

We model the non-ionic interactions with a truncated and shifted Ashbaugh-Hatch potential (***Ashbaugh and Wood, 1997***; ***Ashbaugh and Hatch, 2008***),

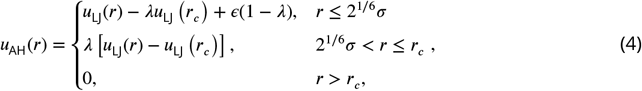

where *ϵ* = 0.8368 kJ mol^−1^, *r*_*c*_ = 2 nm, and the 12-6 Lennard-Jones potential *u*_*LJ*_ is

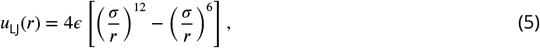

where *σ* is the average of the van der Waals radii of the interacting molecules with values for amino acids determined by Kim & Hummer (***Kim and Hummer, 2008***).

The *λ c*_*ij*_ value for the interactions between particle *i* and *j* is determined as

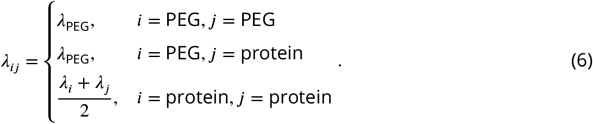

Thus, for protein–protein interactions, *λ* _*ij*_ is the arithmetic average of the amino acid-specific *Jc* values. For protein–PEG and PEG–PEG interactions, we instead use a fixed value of *Jc*_PEG_, which we determined as described below.

### Parameterisation of the PEG model

To parameterise the PEG bead, we determine the values for *λ*_PEG_ and *σ*_PEG_ in a scan within the following ranges: *λ* _PEG_ E {0.0, 0.1, 0.2, 0.3, 0.4} and *σ* _PEG_ ∈ {0.35 nm, 0.40 nm, 0.43 nm, 0.46 nm}. First, we screen the parameters to reproduce the experimental single-chain compaction of PEG with different MWs, which allows us to tune PEG–PEG interactions. Second, we screen the parameters to capture the effect of PEG crowding on IDR chain compaction, as *λ*_PEG_ dictates the PEG–Protein interactions (Eq. 6).

For the first part, we performed simulations of single-chain PEG in a cubic box with simulation settings described in Table 1. We simulate a range of MWs from 400 Da to 30 kDa with all the different values of *λ*_PEG_ and *σ* _PEG_. Subsequently, we calculated the radius of gyration,***R*** _g_, with error estimation through a blocking approach (***Flyvbjerg and Petersen, 1989***) as implemented in the BLOCKING software (github.com/fpesceKU/BLOCKING). We then quantified the agreement with the experimental data by calculating the relative error, (***R***_g,sim._ − ***R***_g,exp._) ***R***_g,exp._ * 100%, and a reduced χ^2^ statistic, 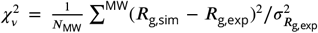, where *N*_MW_ is the total number of data points equal to the number of MWs that we simulated. In this part of the parameterisation, we used an experimental dataset compiled by Sherck and colleagues (***Sherck et al., 2020***). As no errors were reported on the experimental ***R***_g_ values, we assumed a 10% error.

**Table 1.** Settings for single-chain simulations of PEG with different molecular weights.

| MW PEG (Da) | DP | T (K) | L<sub>box</sub> (nm) | R<sub>g,exp.</sub> (nm) | R<sub>ee,exp.</sub> (nm) | Exp.Solv. |
| --- | --- | --- | --- | --- | --- | --- |
| 400 | 9 | 310 | 30 | 0.6 | 1.50 | H<sub>2</sub>O |
| 1000 | 22 | 310 | 30 | 0.9 | 2.10 | H<sub>2</sub>O |
| 1560 | 35 | 333 | 30 | 1.6 | 4.00 | D<sub>2</sub>O |
| 2000 | 45 | 295 | 30 | 1.4 | 3.40 | D<sub>2</sub>O |
| 3000 | 68 | 298 | 30 | 2.2 | 5.50 | D<sub>2</sub>O |
| 4000 | 90 | 295 | 30 | 2.2 | 5.40 | D<sub>2</sub>O |
| 5000 | 113 | 298 | 30 | 3.1 | 7.80 | H<sub>2</sub>O |
| 8000 | 181 | 295 | 35 | 2.6 | 6.50 | D<sub>2</sub>O |
| 10000 | 227 | 298 | 35 | 4.0 | 10.00 | D<sub>2</sub>O |
| 16000 | 363 | 298 | 35 | 5.4 | 13.50 | H<sub>2</sub>O |
| 20000 | 454 | 298 | 40 | 6.1 | 15.25 | D<sub>2</sub>O |
| 21000 | 476 | 333 | 40 | 4.6 | 11.50 | D<sub>2</sub>O |
| 25000 | 567 | 298 | 40 | 6.5 | 16.25 | H<sub>2</sub>O |
| 30000 | 681 | 298 | 50 | 7.5 | 18.75 | D<sub>2</sub>O |

For the second part, where we explored the crowding-induced compaction of IDRs, we simulated a single chain of the proteins ACTR and IN (see Table S1 for sequences) in a cubic box with varying volume fractions of PEG400 and PEG8000, and at the solution conditions detailed by ***Soranno et al***. (***2014***) (Table 2). To reproduce the experimental volume fractions, *ϕ*_PEG_, we calculated the number of PEG chains,*N* _PEG chains_, as

**Table 2.** Simulation settings and experimental data for the single-molecule protein crowding scan with the binding domain of the Nuclear receptor coactivator 3 (ACTR) and the N-terminal domain of HIV-1 integrase (IN) (*Soranno et al., 2014*)

| Protein | MW PEG (Da) | $\phi_{\text{PEG}}$ | $R_g$ (nm) | $R_{ee}$ (nm) | T (K) | $c_s$ (M) |
| --- | --- | --- | --- | --- | --- | --- |
| ACTR | 400 | 0.00 | 2.48 | 5.55 | 295 | 0.11 |
|  |  | 2.51 | 2.44 | 5.45 | 295 | 0.11 |
|  |  | 9.07 | 2.44 | 5.46 | 295 | 0.11 |
|  |  | 14.01 | 2.41 | 5.39 | 295 | 0.11 |
|  |  | 19.17 | 2.36 | 5.27 | 295 | 0.11 |
|  |  | 38.32 | 2.15 | 4.81 | 295 | 0.11 |
|  | 8000 | 0.00 | 2.48 | 5.55 | 295 | 0.11 |
|  |  | 5.68 | 2.40 | 5.36 | 295 | 0.11 |
|  |  | 10.35 | 2.33 | 5.21 | 295 | 0.11 |
|  |  | 13.81 | 2.32 | 5.19 | 295 | 0.11 |
|  |  | 19.90 | 2.28 | 5.10 | 295 | 0.11 |
| IN | 400 | 0.00 | 2.02 | 4.51 | 295 | 0.11 |
|  |  | 3.71 | 2.01 | 4.51 | 295 | 0.11 |
|  |  | 7.78 | 2.01 | 4.49 | 295 | 0.11 |
|  |  | 16.79 | 1.99 | 4.45 | 295 | 0.11 |
|  |  | 25.22 | 1.97 | 4.40 | 295 | 0.11 |
|  |  | 35.97 | 1.95 | 4.37 | 295 | 0.11 |
|  | 8000 | 0.00 | 2.02 | 4.51 | 295 | 0.11 |
|  |  | 4.52 | 2.00 | 4.48 | 295 | 0.11 |
|  |  | 8.12 | 2.00 | 4.47 | 295 | 0.11 |
|  |  | 12.39 | 1.96 | 4.39 | 295 | 0.11 |
|  |  | 17.50 | 1.95 | 4.37 | 295 | 0.11 |

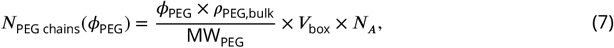

where *ρ*_PEG,bulk_ = 1120 g/L is an average density for bulk low-MW PEG (***Soranno et al., 2014***), MW_PEG_ is the molecular weight of PEG, ***V***_box_ is the volume of the simulation box, and *N*_A_ is Avogadro’s number.

In evaluating the agreement of our model against smFRET experiments, we compared simulated R_ee_ values with R_ee_ values back-calculated from the experimental R_g_ data reported by ***Soranno et al***. (***2014***) using the following relationship: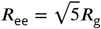. We then calculated a reduced χ^2^ statistic, 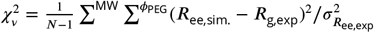, where is the total number of data points.

### Setups for direct-coexistence simulations

We modelled phase separation with direct-coexistence simulations in an elongated rectangular box of dimensions *L*_*z*_ *⪢ L*_*x*_ = *L*. In this manner, we obtain a slab geometry across periodic boundaries (***Blas et al., 2008***; ***Silmore et al., 2017***; ***Regy et al., 2021b***; ***Tesei et al., 2021***). For each IDR, we chose box lengths and numbers of protein chains to obtain a sufficiently thick slab, to minimize finite-size effects, while maintaining a large enough volume of the protein-dilute phase, which is needed to sample *C*_sat_ away from the interface and to prevent slab–slab interactions along the long axis of the box. To reproduce the PEG mass concentrations used in the experiments, we first converted the %w/v values, in g per 100 mL, into *ϕ*_PEG_ using the relationship *ϕ*_PEG_ × ρ_PEG,solid_ = 10 × %w/v, and then calculated the number of PEG chains to include in the simulation box, ρ_PEG chains_, using Eq. 7.

To facilitate equilibration, we initiate simulations with configurations in which proteins and PEG are placed in different subvolumes, separated by *L*_*z*_ 2 along the *z*-axis of the box. The initial topology of each chain is an Archimedean spiral where beads are spaced by the equilibrium bond length. The relevant numerical values of our simulation setups are summarized in Table 3.

**Table 3.** Settings for direct-coexistence simulations of proteins with PEG400 and PEG8000.

| Protein | MW PEG (Da) | [PEG] (%w/v) | $N_{\text{prot.}}$ | $L_{xy}$ (nm) | $L_z$ (nm) | T (K) | $c_s$ (mM) |
| --- | --- | --- | --- | --- | --- | --- | --- |
| hnRNPA1 LCD | 400,8000 | 0.0,2.5,5.0,7.5,10.0 | 100 | 15 | 150 | 298 | 150 |
| Ddx4n1 | 3000 | 0.0,0.7,1.0,1.4,<br>2.0,3.0,4.0 | 100 | 17 | 300 | 293 | 110 |
| αSyn WT | 8000 | 0.0,2.5,5.0,7.5,10.0,<br>12.5,15.0,20.0 | 100 | 15 | 150 | 298 | 150 |
| αSyn Swap Variant | 8000 | 0.0,2.5,5.0,7.5,10.0 | 100 | 15 | 150 | 298 | 150 |

### Analysis of direct-coexistence simulations

We quantify the propensity of a protein to phase separate as the transfer free energy,

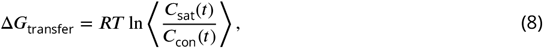

where is the gas constant whereas *C*_sat_(*t*) and *C*_con_(*t*) are the time series of the protein concentrations in the protein-dilute and -dense phases, respectively.

To calculate the concentrations in the protein-dense, C_con_, and protein-dilute, *C*_sat_, phases, we align the protein slab to the centre of the simulation box and determine the boundaries between the phases based on the equilibrium concentration profile of the protein as described in previous work (***Jung and Yethiraj, 2018***; ***Tesei and Lindorff-Larsen, 2022***; ***von Bülow et al., 2024***). The boundaries along the *z*-axis of the box used in this study are reported in the DataFrames deposited in the GitHub repository supporting the analyses of this work.

## Data and Code Availabillity

All the data and code used for this work is available via https://github.com/KULL-Centre/_2025_rauh_peg. The PEG model is available in the CALVADOS package https://github.com/KULL-Centre/CALVADOS.

## Acknowledgements

We thank members of SBiNLab and Tanja Mittag and Rohit Pappu for helpful discussions. We acknowledge access to computational resources from the Biocomputing Core Facility at the Department of Biology, University of Copenhagen, from the Resource for Biomolecular Simulations (ROBUST; supported by the Novo Nordisk Foundation; NNF18OC0032608), and the Danish National Supercomputer for Life Sciences (Computerome). This work is a contribution from the PRISM (Protein Interactions and Stability in Medicine and Genomics) centre funded by the Novo Nordisk Foundation (to K.L.-L.; NNF18OC0033950).

