## Supporting Figures and Tables for "A coarse-grained model for disordered proteins under crowded conditions"

### Supplemental tables

| Protein | Sequence | source |
|---|---|---|
| **ACTR** | GTQNRPLLRNSLDDLVGPPSNLEGQSDERALLDQLHTLLSNTDATGLEEI DRALGIPELVNQGQALEPKQD | (***Soranno et al., 2014***) |
| **IN** | CAQEEHEKAHSNFRAMASDFNLPPVVAKEIVASCDKCQLKGEAMHGQVD | (***Soranno et al., 2014***) |
| **ProT$\alpha$-C** | CEEGGEEEEEEEEGDGEEEDGDEDEEAESATGKRAAEDDEDDDVDTKKQK TDED | (***Soranno et al., 2014***) |
| **ProT$\alpha$** | SDAAVDTSSEITTKDLKEKKEVVEEAENGRDAPANGNAENEENGEQEADN EVDEECEEGGEEEEEEEEGDGEEEDGDEDEEAESATGKRAAEDDEDDDVD TKKQKTDEDC | (***Soranno et al., 2014***) |
| **A1-LCD WT** | GSMASASSSQRGRSGSGNFGGGRGGGFGGNDNFGRGGNFSGRGGFGGSRG GGGYGGSGDGYNGFGNDGSNFGGGGSYNDFGNYNNQSSNFGPMKGGNFGG RSSGGSGGGGQYFAKPRNQGGYGGSSSSSSYGSGRRF | (***Martin et al., 2020***) |
| **A1-LCD Aro$^{-}$** | GSMASASSSQRGRSGSGNSGGGRGGGFGGNDNFGRGGNSSGRGGFGGSRG GGGYGGSGDGYNGFGNDGSNSGGGGSSNDFGNYNNQSSNFGPMKGGNFGG RSSGGSGGGGQYSAKPRNQGGYGGSSSSSSSGSGRRF | (***Martin et al., 2020***) |
| **A1-LCD Aro$^{--}$** | GSMASASSSQRGRSGSGNSGGGRGGGFGGNDNSGRGGNSSGRGGFGGSRG GGGSGGSGDGYNGSGNDGSNSGGGGSSNDFGNSNNQSSNSGPMKGGNFGG RSSGGSGGGGQYSAKPRNQGGSGGSSSSSSSGSGRRS | (***Martin et al., 2020***) |
| **Ddx4n1** | MGDEDWEAEINPHMSSYVPIFEKDRYSGENGDNFNRTPASSSEMDDGPSR RDHFMKSGFASGRNFGNRDAGECNKRDNTSTMGGFGVGKSFGNRGFSNSR FEDGDSSGFWRESSNDCEDNPTRNRGFSKRGGYRDGNNSEASGPYRRGGR GSFRGCRGGFGLGSPNNDLDPDECMQRTGGLFGSRRPVLSGTGNGDTSQS RSGSGSERGGYKGLNEEVITGSGKNSWKSEAEGGES | (***Stender et al., 2021***) |
| $\alpha$**Syn** | MDVFMKGLSKAKEGVVAAAEKTKQGVAEAAGKTKEGVLYVGSKTKEGVVH GVATVAEKTKEQVTNVGGAVVTGVTAVAQKTVEGAGSIAAATGFVKKDQL GKNEEGAPQEGILEDMPVDPDNEAYEMPSEEGYQDYEPEA | (***Dedmon et al., 2005***) |
| $\alpha$**Syn Swap Variant** | MGMAPKKKAKKGIMGKKGTLKKKKKYSTKIAAVPGQVKGGTTTVNPVVVV GGVQYALVQPGGSDVVEAMKTVTAVVNGEGGESEAEQEAQVTALVVAFPQ ANSTVTHAEGGFLAGDADEGYEEDDDEEYEEEEEVEAAAA | (***Pesce et al., 2024***) |

**Table S1.** Sequences of proteins simulated in this study.

| Protein | $f_K$ | $f_R$ | $f_E$ | $f_D$ | $f_{ARO}$ | FCR | $\langle\lambda\rangle$ | SHD | SCD | $\kappa$ | NCPR |
|---|---|---|---|---|---|---|---|---|---|---|---|
| **ACTR** | 0.014 | 0.056 | 0.085 | 0.099 | 0.0 | 0.413 | 0.268 | 3.183 | 0.584 | 0.155 | -0.099 |
| **IN** | 0.082 | 0.02 | 0.102 | 0.061 | 0.041 | 0.359 | 0.286 | 2.517 | -0.063 | 0.109 | -0.041 |
| **ProT$\alpha$-C** | 0.074 | 0.019 | 0.37 | 0.222 | 0.0 | 0.186 | 0.704 | 1.324 | 21.879 | 0.528 | -0.481 |
| **ProT$\alpha$** | 0.073 | 0.018 | 0.309 | 0.164 | 0.0 | 0.22 | 0.573 | 1.87 | 34.939 | 0.405 | -0.373 |
| **A1-LCD WT** | 0.015 | 0.073 | 0.0 | 0.029 | 0.139 | 0.611 | 0.124 | 5.506 | 1.796 | 0.194 | 0.066 |
| **A1-LCD Aro$^-$** | 0.015 | 0.073 | 0.0 | 0.029 | 0.095 | 0.592 | 0.124 | 5.331 | 1.796 | 0.194 | 0.066 |
| **A1-LCD Aro$^{--}$** | 0.015 | 0.073 | 0.0 | 0.029 | 0.044 | 0.569 | 0.124 | 5.127 | 1.796 | 0.194 | 0.066 |
| **Ddx4n1** | 0.034 | 0.102 | 0.076 | 0.076 | 0.093 | 0.488 | 0.292 | 4.962 | -0.948 | 0.230 | -0.013 |
| **$\alpha$Syn** | 0.107 | 0.0 | 0.129 | 0.043 | 0.043 | 0.341 | 0.286 | 3.081 | -2.076 | 0.169 | -0.057 |
| **$\alpha$Syn Swap Var.** | 0.107 | 0.0 | 0.129 | 0.043 | 0.043 | 0.341 | 0.286 | 3.099 | -13.796 | 0.592 | -0.057 |

**Table S2.** Descriptors of the sequence composition and patterning for proteins simulated in these studies. For charged residue composition and patterning: $f_K$, $f_R$, $f_E$, $f_D$, FCR, SCD & $\kappa$. For hydrophobic & aromatic residue composition and patterning (***Das and Pappu, 2013***; ***Sawle and Ghosh, 2015***; ***Zheng et al., 2020***).

| Ddx4n1 | Dilute Phase Conc. (μM) | | | |
|---|---|---|---|---|
| $\phi_{PEG3000}$ | Set 1 | Set 2 | Average | SD |
| 2 | 49 | 50 | 49 | 0.4 |
| 3 | 38 | 39 | 38 | 1 |
| 4 | 33 | 34 | 33 | 0.4 |
| 5 | 27 | 28 | 28 | 1 |
| 6 | 23 | 24 | 24 | 0.4 |
| 7 | 18 | 18 | 18 | 0.0 |

**Table S3.** Experimental $C_{sat}$ data for Ddx4n1 with increasing PEG3000 concentrations (***Stender et al., 2021***).

| Ddx4n1 | Dilute Phase Conc. (μM) | | | |
|---|---|---|---|---|
| Overall Conc. (μM) | Set 1 | Set 2 | Average | SD |
| 100 | 95 | 100 | 99 | 3 |
| 116 | 97 | 107 | 103 | 5 |
| 133 | 92 | 100 | 97 | 4 |
| 167 | 96 | 105 | 102 | 5 |

**Table S4.** Experimental $C_{sat}$ data for Ddx4n1 measured with varying protein concentrations (***Stender et al., 2021***).

9 ## Supplemental figures

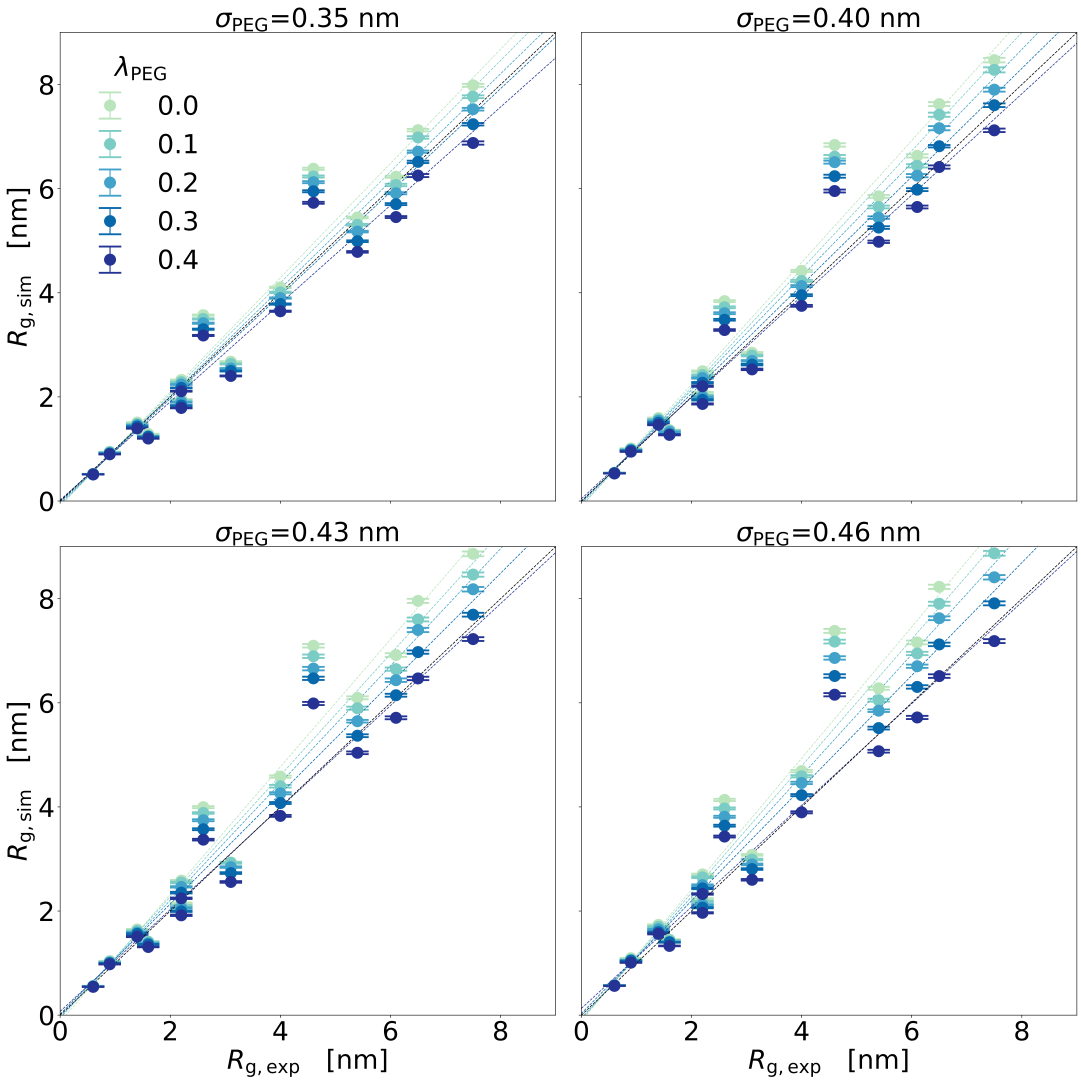


10

**Figure S1.** Comparison of single chain PEG compaction from simulations and experiments.

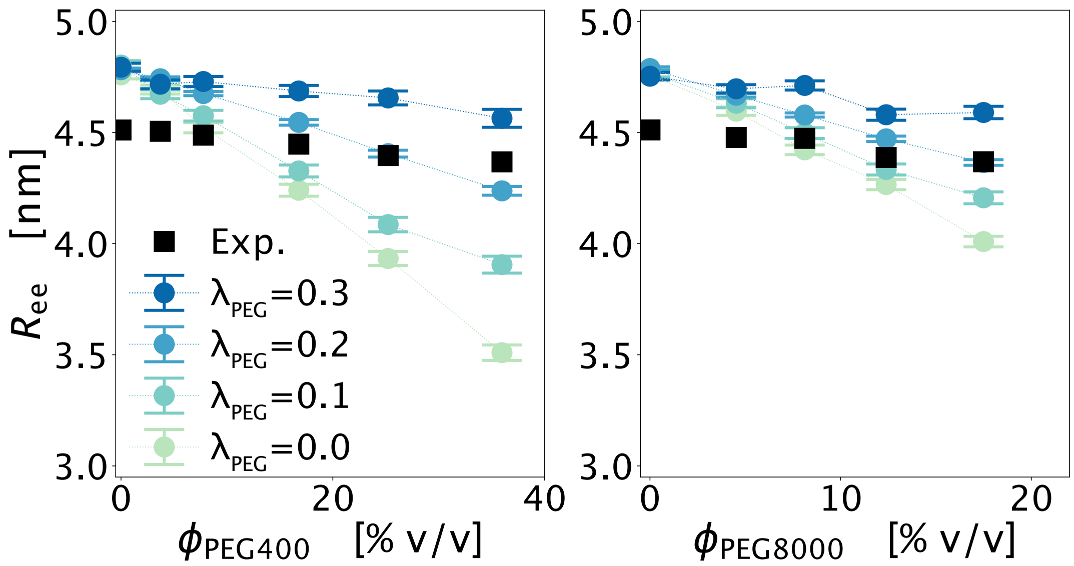


11

**Figure S2.** Protein compaction as a function of PEG volume fractions for the N-terminal domain of HIV-1 integrase (IN)

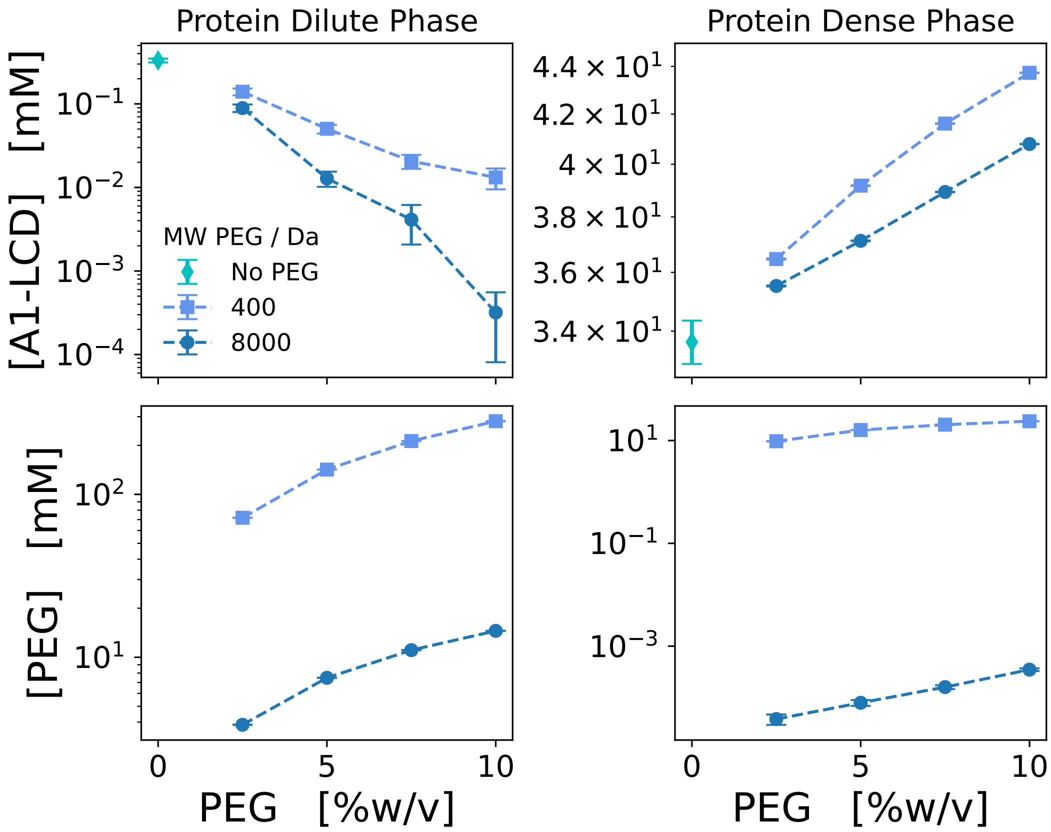


**Figure S3.** Concentrations of A1 and PEG in the protein (right panels) and dense (left panels) phases for A1 with PEG400 (square) and PEG8000 (dot) plotted against the PEG concentration in %w/v.

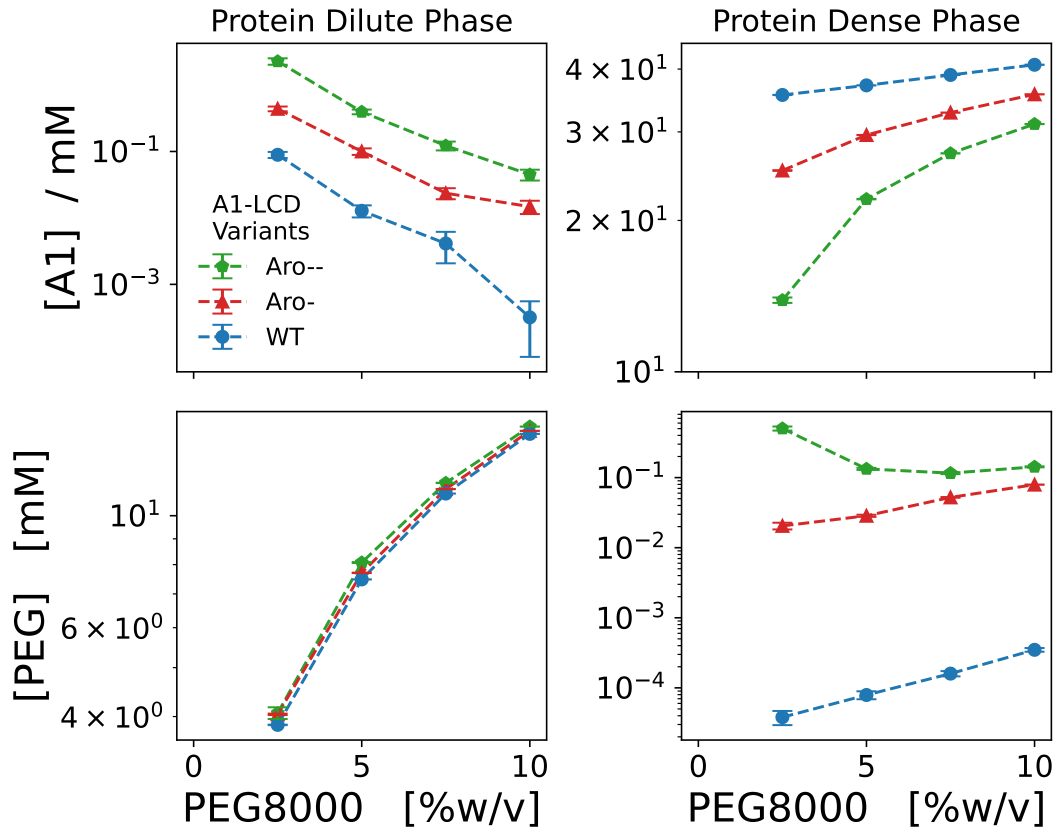


**Figure S4.** Concentrations of protein in the protein (right panels) and dense (left panels) phases for the WT hnRNPA1-LCD (blue dots) and the aromatic variants Aro$^{-}$ (red triangles) and Aro$^{--}$ (green pentagons) plotted against the PEG concentration in %w/v

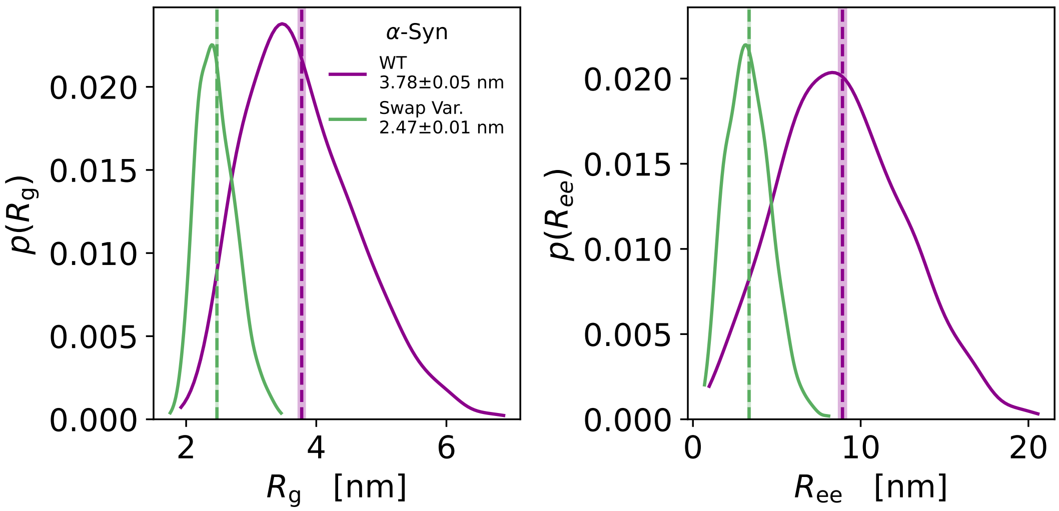


14

**Figure S5.** Distribution of the radius of gyration $R_g$ and end-to-end distance $R_{ee}$ for WT $\alpha$-synuclein (dark magenta) and for the swap variant of $\alpha$-synuclein (green) sampled from single chain CALVADOS 2 simulations.

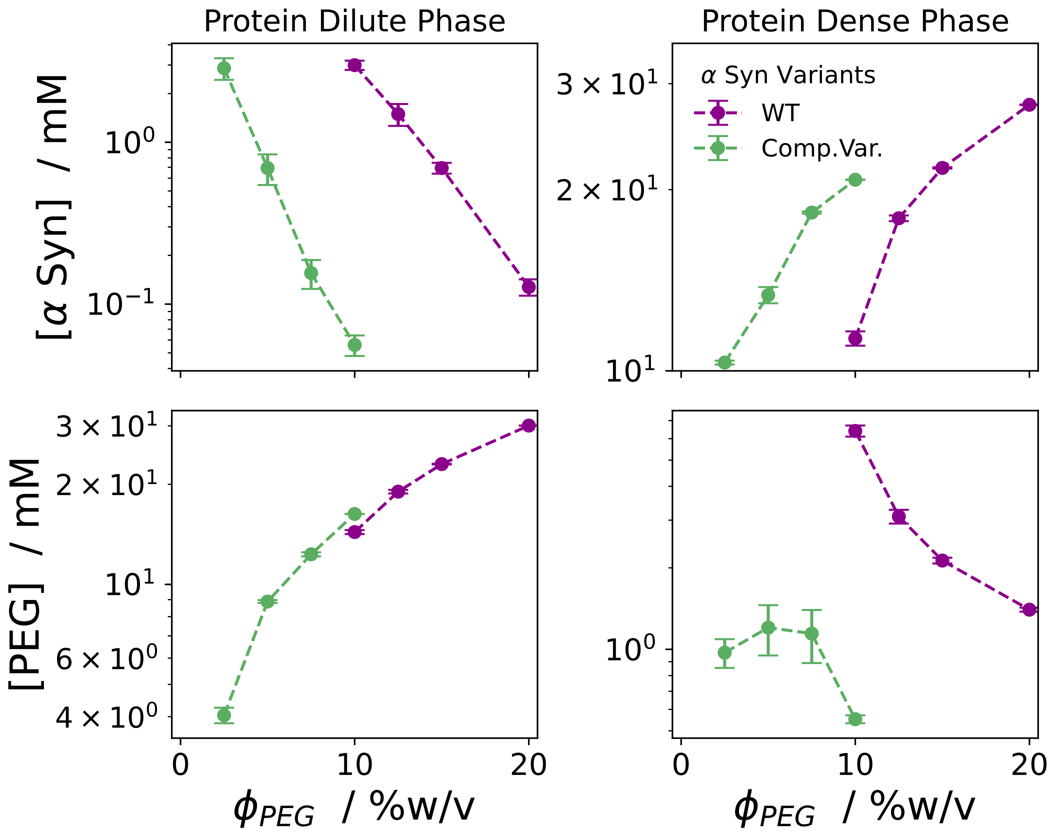


**Figure S6.** Concentrations of protein and PEG8000 in the protein dilute (right panels) and dense (left panels) phases as a function of the overall PEG concentration for WT αSyn (dark magenta) and the swap variant (green).

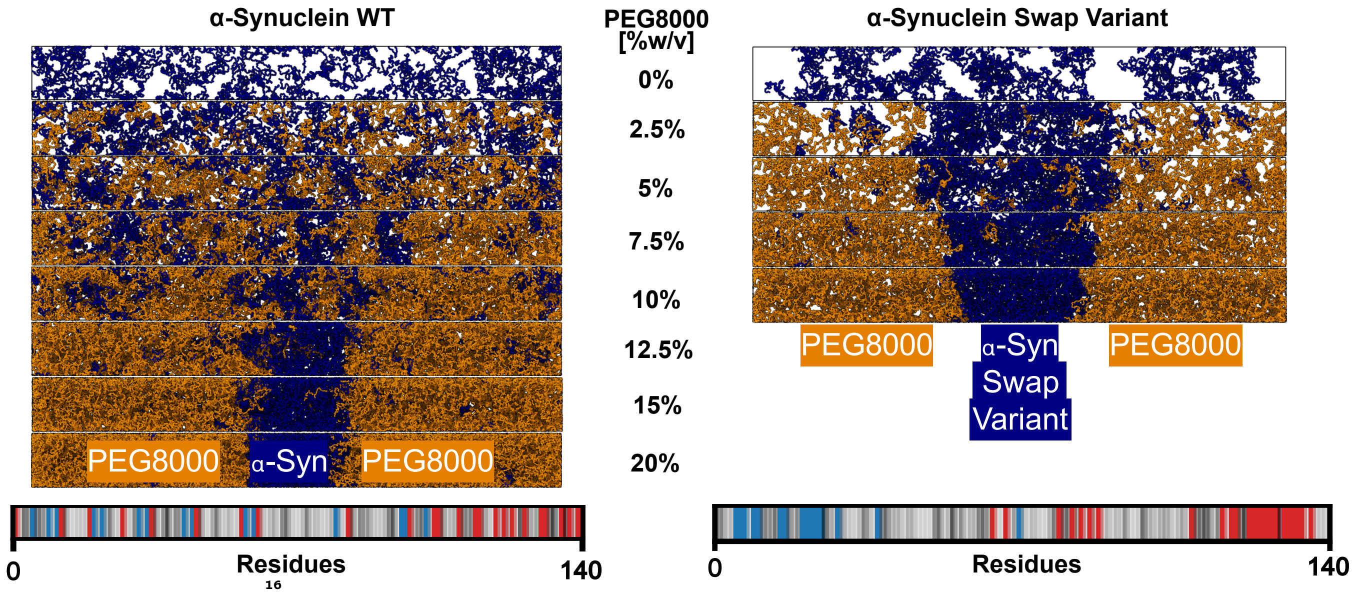


**Figure S7.** On top, representative snapshots of the different titration stages for WT $\alpha$Syn and the swap variant at the different PEG8000 concentrations. On the bottom a graphical representation of the sequences, to visualise the difference in charge-patterning between the the wildtype and swap variant. Negative and positive charges are coloured in red and blue, respectively. The neutral residues are coloured by a gray scale that reflects their hydrophobicity (corresponding to the $\lambda$ parameter in CALVADOS).
